# TiSAM: fusing spatial and histopathology foundation models for whole-slide image segmentation

**DOI:** 10.64898/2026.10.16.759688

**Authors:** Zhiyuan Ding, Lucie Dequiedt, Marta Forjaz, Valentina Matos-Romero, Yu Shen, Talia Benducci, André Forjaz, Xiaoke Fei, Yutong Zhu, Patricia Nwoga, William Dhana, Julie Pascalin, Sam S. Lee, Luis Garza, Seung Hyun Lee, Edward J. Pearce, Marcia Irene Canto, Jennifer P. Morton, Daniela Cihakova, Denis Wirtz, Data Science TeamLab, Ashley L. Kiemen

**Author notes:** Send Correspondence to: Ashley L. Kiemen Department of Pathology, 1729 E Monument St, Office 10219, Baltimore, MD 21287. indicates co-first authorship (Zhiyuan Ding, Lucie Dequiedt).

## Abstract

Semantic segmentation of whole-slide images (WSIs) quantifies tissue composition and architecture, but existing models require extensive annotation and lose accuracy as the number of tissue classes increases or on heterogeneous, cross-scanner datasets. Here, we present the Tissue Segment Anything Model (TiSAM), which fuses SAM3 spatial features with UNI2-h pathology features within a multi-scale decoder. Because public benchmarks contain few tissue classes, we release MegaMouse, comprising 36,833 annotations of 52 tissue classes from 12 mouse organs, and an 11-class human pancreas dataset. TiSAM achieved the highest overall performance across four public datasets compared to existing convolutional neural network and vision transformer models, maintained high accuracy on MegaMouse, and maintained high accuracy in the human pancreas dataset when trained on as few as 5 annotated WSIs where previous integrations of SAM with pathology foundation models did not. TiSAM segmentations were also more consistent across eight slide scanners compared to other models. These results establish TiSAM as a general and highly adaptable framework for tissue segmentation, enabling studies containing many tissue classes, limited annotations, or images from multiple scanners.

## Introduction

Semantic segmentation of whole slide images (WSIs) enables quantitative measurements of tissue composition and architecture at a scale difficult to achieve through manual annotation and is useful for identifying spatial niches for downstream molecular profiling^1–4^. Early segmentation approaches in pathology relied largely on convolutional neural networks (CNNs), which have since been widely adopted for diverse segmentation tasks^5–8^. Among CNN-based frameworks, DeepLabV3+ with ResNet-50 backbones has demonstrated robust performance^9,10^. However, CNNs have been shown to have limited generalizability across tissues, staining protocols, and cohorts, and in tasks with high numbers of tissue classes, requiring substantial annotated data for fine-tuning^11,12^.

Vision transformer (ViT)-based architectures have expanded image segmentation beyond task-specific CNNs. The Segment Anything Model (SAM) introduced a generalist, promptable framework capable of zero-shot and few-shot segmentation across diverse image domains^13^. Subsequent versions, SAM2 and SAM3, and adaptations to medical imaging have further extended its capabilities^14–17^. However, SAM and its successors lack domain-specific knowledge of tissue morphology, limiting their performance on histopathology segmentation. PathSegmentor^18^ recently extended this paradigm by integrating semantic segmentation via language prompts. In parallel, pathology foundation models have emerged that are specifically pretrained on large collections of histopathology images using self-supervised learning^19–21^. Unlike segmentation foundation models, they serve as general-purpose feature encoders and require a separate decoder to produce pixel-level segmentation.

Recent studies have combined SAM with pathology foundation models for prompt-free semantic segmentation^22,23^. However, these approaches adapt SAM through learned prompts or augmented inputs, with features from both encoders concatenated at a single spatial scale and decoded using one learned prompt per class. Evaluation has been limited to public image tiles with five or fewer classes. It therefore remains unclear whether these models outperform CNN-based architectures with many tissue classes, limited training annotations, or tissue heterogeneity stemming from differences in slide scanner.

Here, we present the Tissue Segment Anything Model (TiSAM), a framework that fuses pathology-specific semantic and SAM-derived spatial features within a multi-scale decoder for efficient WSI segmentation while preserving fine-grained tissue boundaries. We compare TiSAM with eight existing model architectures across four tasks: (1) benchmarking on public challenge datasets; (2) learning a large number of classes; (3) learning from minimal training annotations and (4) segmentation consistency across scanners. TiSAM ranked among the strongest-performing models across all four public datasets. To evaluate performance with many classes and limited annotations, we establish two publicly available benchmarking datasets; an 11-class human pancreas dataset comprising clinically diverse WSIs and MegaMouse, a 52-class dataset spanning 12 mouse organs. On MegaMouse, TiSAM achieved 88.6% class-balanced accuracy, compared with 45% for SAM-Path, 18.2% for SAM2-Path, and 1.5% for pretrained PathSegmentor. When trained on only 5 or 10 pancreas WSIs, TiSAM achieved macro-F1 scores of 93.3% and 93.5%, respectively, compared with 86.1% and 91.0% for DeepLabV3+. Across eight slide scanners, TiSAM segmentations were more consistent than those of DeepLabV3+, SAM-Path, SAM2-Path, and PathSegmentor. These results show that previous integrations of SAM with pathology foundation models do not consistently outperform lightweight CNN-based architectures, whereas TiSAM’s multi-scale fusion maintains strong performance across diverse tasks. Together, these results establish TiSAM as a general framework for WSI tissue segmentation, with utility for tasks involving many tissue classes, limited annotations, or multiple scanners.

## Results

### TiSAM fuses SAM3 and UNI2 using an optimized multi-scale decoder

To determine the performance of fused representations from histopathology and general-purpose vision foundation models in WSI tissue segmentation, we developed TiSAM, which combines a frozen histopathology foundation model with a SAM encoder **(Fig. 1A)**. The pathology foundation model extracts semantic representations learned from large-scale pathology datasets, whereas SAM extracts spatial representations that support precise localization and boundary delineation. Pathology features are projected into the 256-dimensional SAM3 feature space and integrated with unprojected SAM3 features through a multi-scale fusion module, after which a pixel decoder aggregates features across spatial resolutions to generate high-resolution spatial embeddings. These representations are then passed to a query-based mask decoder adapted from Mask2Former^24^. The decoder uses learnable queries to selectively aggregate information from the fused feature maps through self-and cross-attention, generating mask embeddings that are compared with spatial pixel embeddings to produce dense semantic masks. This design enables the decoder to integrate semantic and spatial information while preserving the high-resolution spatial structure required for whole-slide tissue segmentation **(Fig. S1)**.

**Figure 1:**
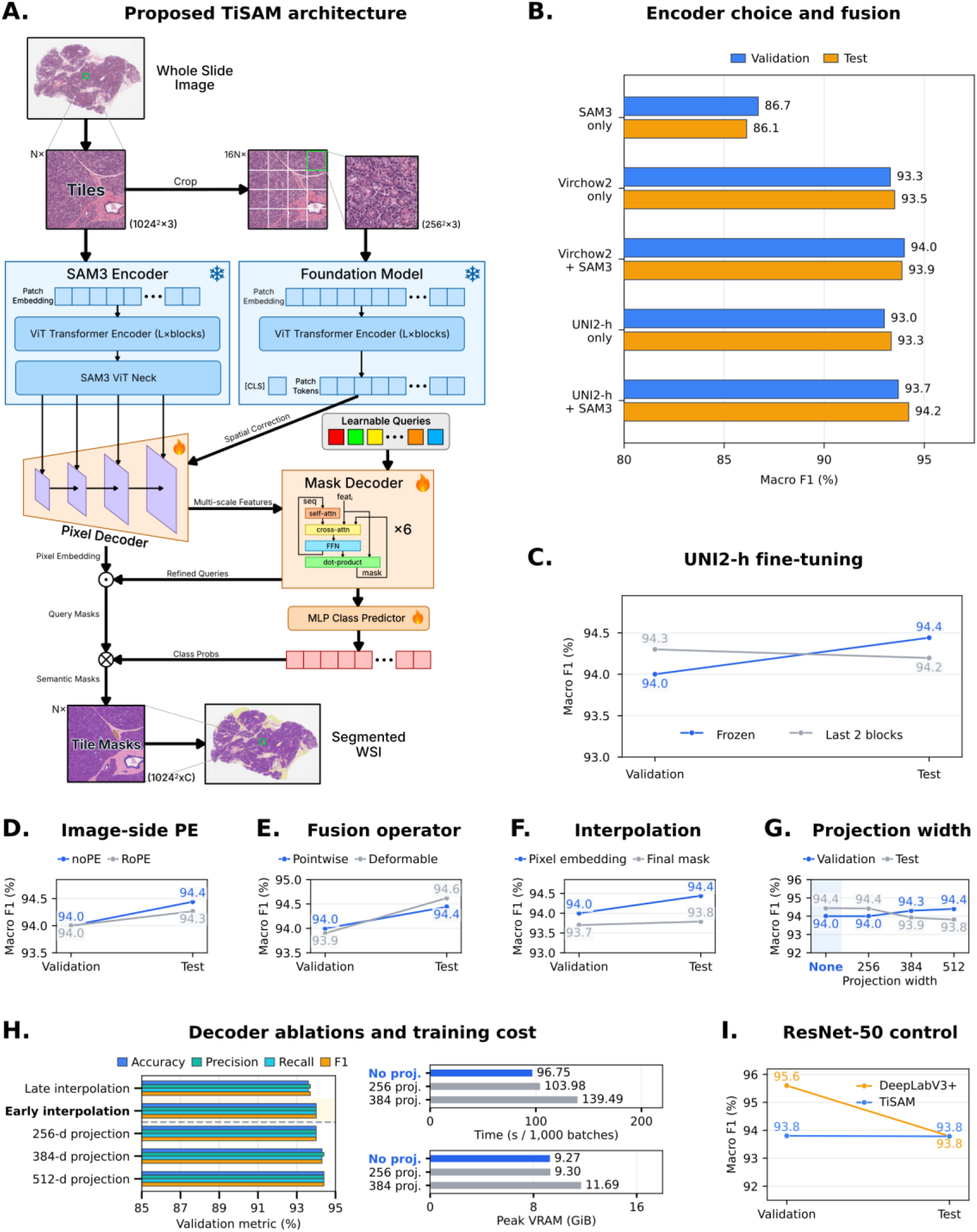
TiSAM fuses SAM3 and UNI2 using an optimized multi-scale decoder. **A.** TiSAM combines frozen SAM3 and pathology foundation model encoders with trainable pixel and mask decoders to generate whole-slide semantic segmentation. Snowflakes and flames indicate frozen and trainable components. **B.** Comparison of Virchow2 and UNI2-h, alone and combined with SAM3, and SAM3 alone. **C.** Frozen UNI2-h versus fine-tuning its final two blocks. **D.** Image-side positional encoding: no positional embedding (noPE) versus rotary positional embedding (RoPE). **E.** Pointwise versus deformable feature fusion. **F.** Interpolation of pixel embeddings versus final masks. **G.** Effect of feature projection width. **H.** Decoder ablations comparing interpolation strategies and feature projection widths, with validation metrics, training time, and peak GPU memory. Measurements were obtained on an NVIDIA RTX PRO 6000 GPU using a batch size of 1 and gradient accumulation over 16 batches (effective batch size, 16). Training time was measured end to end, including encoder computation, and is reported per 1,000 batches. **I.** TiSAM versus DeepLabV3+ with a ResNet-50 encoder. Panels **B-G** and **I** show validation and test foreground macro-averaged F1.

Architectural ablation experiments in this section were conducted on a human pancreas dataset. We first evaluated whether both feature streams were necessary by systematically ablating the architecture and comparing models on an internal pancreas segmentation dataset using either encoder alone or their combination **(Fig. 1B).** We evaluated two popular pathology foundation models, Virchow2^25^ and UNI2-h^21^, with and without the SAM3 encoder **(Fig. 1B)**. When used alone, both foundation models performed similarly, with a validation F1 score of approximately 93%, with Virchow2 showing a modest advantage on the held-out test set. Using the SAM3 encoder without histopathology features resulted in substantially lower performance, with a validation F1 score of approximately 87%. Combining the SAM3 encoder with either UNI2-h or Virchow2 improved test performance, with the UNI2-h + SAM3 combination achieving the highest test F1. We therefore selected UNI2-h as the pathology foundation model for the final TiSAM architecture, though we note that the modular nature of the TiSAM architecture makes it flexible to future adaptation to better performing pathology foundation model encoders. Further tests showed that fine-tuning the last two blocks of the UNI2-h model modestly improved validation F1 but did not improve test F1 **(Fig. 1C)**.

Finally, we optimized how these representations were integrated within TiSAM. We examined different architectural choices, including image-level positional encoding, the feature-fusion operator, and interpolation strategy **(Fig. 1D-F)**. Changes to positional encoding and feature fusion produced only modest differences in performance, with the default configuration remaining competitive across validation and test sets. Early interpolation, in which feature representations were aligned before mask generation, consistently outperformed later interpolation of the predicted masks. We then evaluated the effect of an additional SAM3 feature projection and its width **(Fig. 1G)**. Increasing the width to 512 channels yielded the highest validation F1, but wider projections reduced test F1, suggesting that the additional capacity did not improve generalization. The no-projection configuration, which retained the projection of pathology features into the 256-dimensional SAM3 feature space but omitted an additional projection of SAM3 features, matched the 256-channel projection variant on both splits. It also reduced computational cost, requiring approximately 97 s per 1,000 training batches, compared with 104 s for the 256-channel projection and 140 s for the 384-channel projection, with corresponding peak training memory usage of 9.3, 9.3, and 11.7 GiB **(Fig. 1H)**. We therefore selected the no-projection configuration for the final TiSAM architecture, retaining pathology feature alignment while avoiding the additional SAM3 projection computation.

To assess the TiSAM decoding framework independently of the foundation-model encoders, we replaced both pretrained encoders with a fully fine-tuned ImageNet-initialized ResNet-50 and compared the resulting TiSAM decoder with a matched DeepLabV3+ decoder using the same ResNet-50 backbone **(Fig. 1I)**. Although DeepLabV3+ achieved higher class-balanced macro-F1 on the validation set (95.6% versus 93.8%), the two decoders achieved essentially identical performance on the held-out test set (both 93.8%), demonstrating that the TiSAM decoder remains competitive beyond the specific foundation-model encoders.

For the remainder of this manuscript, we use TiSAM to refer to the TiSAM model design described above, using UNI2-h as the pathology foundation model encoder. We refer to this version in figures as TiSAM (UNI2-h) to differentiate it from a variation used in benchmarking experiments, TiSAM (ResNet-50). Further, we have integrated TiSAM into CODAvision^9^, a user-friendly interface for annotation generation, model training, and inference **(Fig. S8).**

### TiSAM achieves state-of-the-art performance on public tissue segmentation datasets

Next, we evaluated the performance and generalizability of TiSAM across four publicly available histopathology segmentation datasets spanning breast, lung, and skin tissue, primary and metastatic cancer, and stemming from several different institutions (**Fig. 2A**). The Breast Cancer Semantic Segmentation (BCSS) dataset contains more than 20,000 tissue-region annotations within representative regions of interest selected from 151 breast cancer whole-slide images in The Cancer Genome Atlas (TCGA)^26^. IGNITE H&E comprises annotated tiles from 155 patients with non-small cell lung cancer for multiclass segmentation of primary and metastatic disease, with data collected at multiple centers and using multiple scanners^27^. PUMA comprises 155 primary and 155 metastatic melanoma H&E-tiles with expert-verified tissue and nucleus-level annotations^28^. Finally, BEETLE comprises tiles from 587 breast cancer cases for training and an independent test set comprising 170 densely annotated tiles from 54 cases^29^.

**Figure 2:**
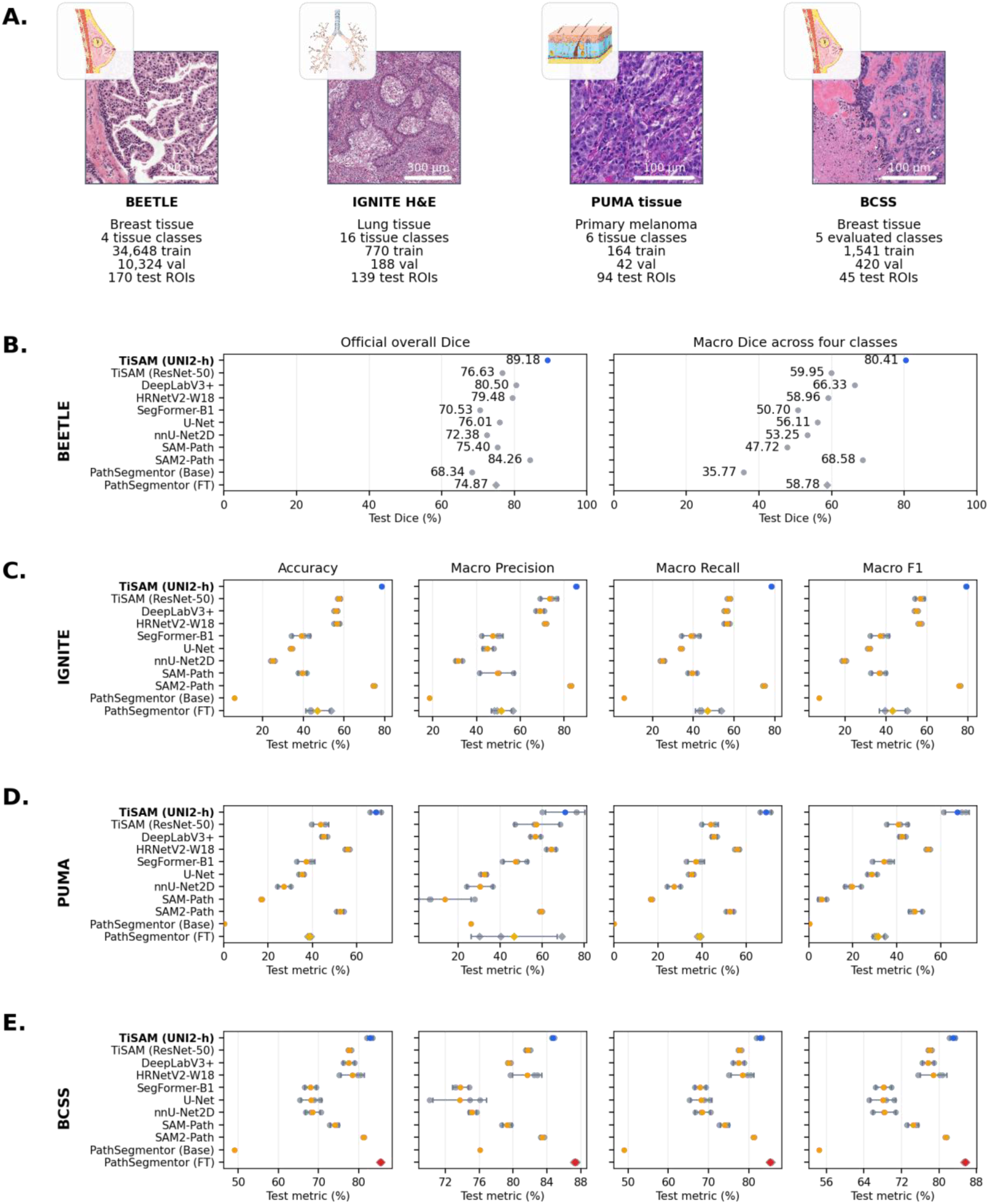
TiSAM achieves state-of-the-art performance on public tissue segmentation datasets. **A.** Representative tissue images, class counts, and dataset sizes for BCSS, IGNITE H&E, PUMA, and BEETLE. Organ illustrations are adapted from Servier Medical Art (CC BY 4.0). **B.** BEETLE test performance measured by official overall Dice and macro-Dice across four classes, including Other, for one validation-selected submission per model. **C–E.** Comparison of TiSAM and baseline models on IGNITE test, PUMA test, and BCSS test, respectively, using accuracy and macro-averaged precision, recall, and F1. Colored markers indicate means, gray markers show individual seeds, and error bars represent sample standard deviations. Comparisons are made within each dataset; horizontal scales vary.

We compared TiSAM with five commonly used segmentation architectures (DeepLabV3+, HRNetV2-W18^30^, SegFormer-B1^31^, U-Net^32^ and nnU-Net2D^33^), two SAM-based pathology models (SAM-Path^22^ and SAM2-Path^23^) and PathSegmentor^18^, a pathology segmentation foundation model that enables natural-language-guided segmentation. We also evaluated a TiSAM variant using a ResNet-50 encoder instead of UNI2-h and SAM3 to assess the contribution of the two foundation models. Performance was evaluated using pixel-level accuracy and macro-averaged precision, recall and F1, calculated from the aggregate confusion matrix for each dataset. Across the independent test sets, TiSAM consistently ranked among the strongest-performing models. TiSAM achieved the highest score across all metrics on IGNITE H&E and PUMA, and the highest overall macro-dice scores on the external BEETLE test set **(Fig. 2B-E)**. Notably, the largest performance advantage of TiSAM over competing architectures was observed on IGNITE H&E and BEETLE: IGNITE comprises 16 tissue classes, providing a challenging test set of multi-class segmentation, while BEETLE includes multiple scanners and sites and an independent external test set, providing a stringent assessment of generalization across acquisition domains. On BCSS, TiSAM achieved the second highest accuracy, recall, and F1, behind a fine-tuned PathSegmentor. This is expected given that BCSS was already included in the original training data of PathSegmentor. Further, while we finetune the weights of PathSegmentor here to make the most comprehensive comparison test, we note that the model is intended for text-prompted segmentation using its pretrained weights, shown in the figure as PathSegmentor (Base), which we consistently outperform.

Together, these results demonstrate that TiSAM consistently outperforms both conventional segmentation architectures and existing SAM-pathology foundation model integrations across independent datasets spanning diverse tissues, diseases, and acquisition settings. Across the four datasets, the consistent performance of TiSAM supports the generalizability of combining SAM3 spatial representations with the pathology-specific semantic representations of UNI2-h.

### TiSAM outperforms existing models in complex multiclass segmentation tasks

Although the publicly available datasets provided a broad assessment of TiSAM’s ability to generalize across tissues, disease states and imaging settings, existing publicly available benchmarking datasets contain a limited number of tissue classes. As a result, these benchmarks did not fully challenge TiSAM’s ability to distinguish between numerous closely related structures. To evaluate this, we developed MegaMouse **(Fig. 3A)**, a novel dataset comprising 36,833 expert-informed manual annotations across 120 WSIs and 12 adult mouse organs: lung, pancreas, skin, kidney, liver, bladder, heart, colon, vagina, uterus, fallopian tube, and ovary. This dataset fuses tissue from diverse disease models, including projects studying cancer, tissue regeneration, infection, inflammation, and aging^34–38^. We fused common structures that existed in multiple organs, such as vasculature, nerves, and adipose tissue, into single, pan-organ classes, and added separate classes for each additional organ-specific structure, such as renal tubules and ovarian follicles, resulting in a total of 52 learnable classes. We finetuned the same group of architectures used previously and compared performance on a held-out test set. TiSAM achieved the highest performance across class-balanced accuracy, macro-precision, macro-recall and macro-F1 **(Fig. 3B)**. To examine class-specific performance, we compared precision, recall and F1 between TiSAM and DeepLabV3+ across all 52 classes **(Fig. 3C, D)**. TiSAM achieved higher precision in 28 classes, compared with 23 for DeepLabV3+ and one tie, and higher recall in 33 classes, compared with 19 for DeepLabV3+. The largest precision gains were observed for esophagus (+21.5 p.p.), collagen (+15.3 p.p.) and ovarian cyst (+10.6 p.p.), whereas the largest decreases occurred for pancreatic precancer (-13.0 p.p.), epidermis (-8.7 p.p.) and normal pancreatic duct (-8.1 p.p.). For recall, TiSAM showed the largest gains for germinal epithelium (+34.7 p.p.), ventricle (+34.2 p.p.) and granuloma (+30.3 p.p.) compared with the largest decreases for immune aggregate (-21.9 p.p.), metaplasia (-12.0 p.p.) and hair follicle (-10.6 p.p.). Overall, TiSAM achieved higher F1 scores in 34 of 52 classes. TiSAM’s improvements were distributed over a wide range of organs, with its largest gains well exceeding its largest losses in both precision and recall.

**Figure 3.**
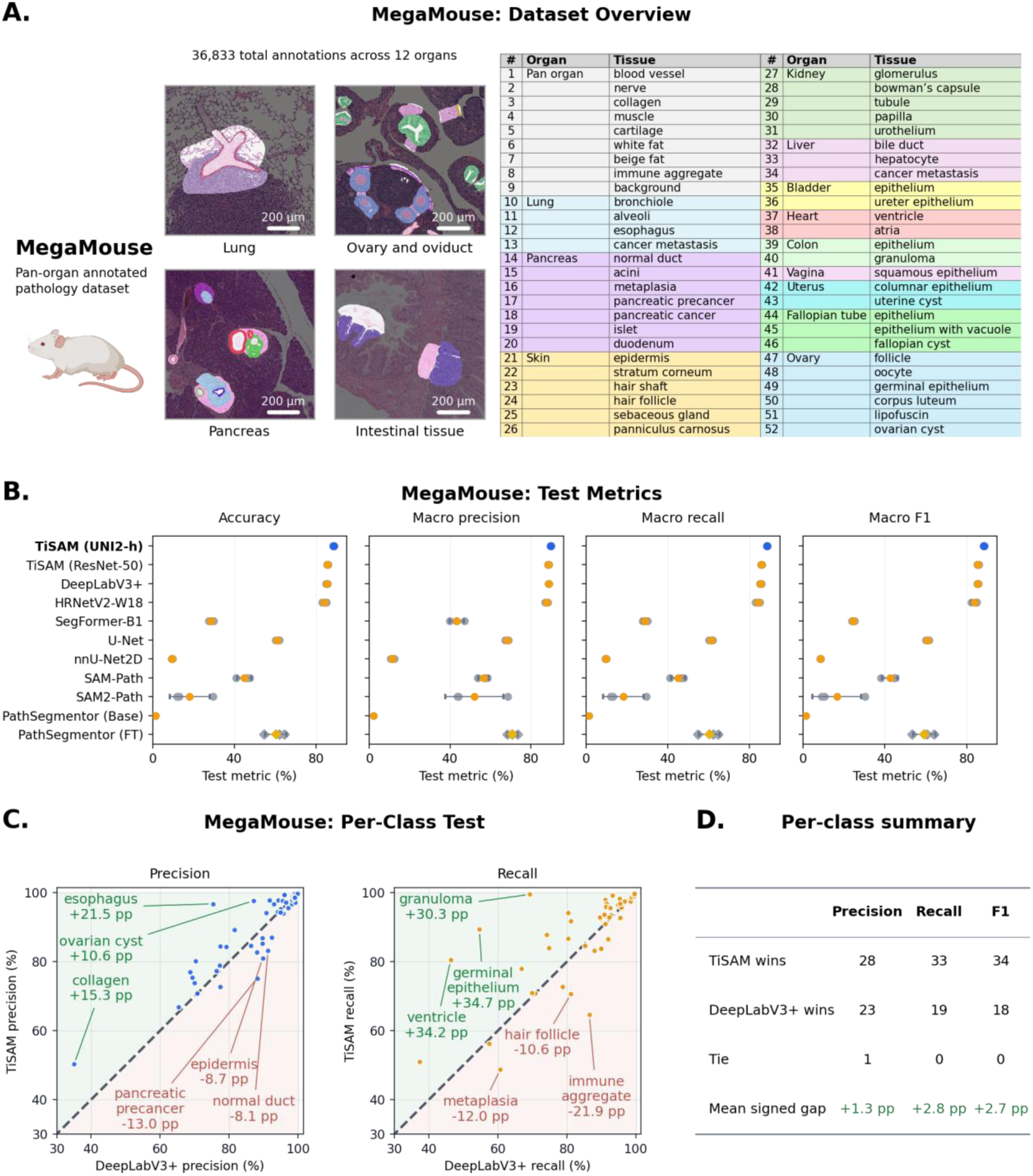
MegaMouse dataset overview and test performance across 52 tissue classes. **A.** Representative annotated tissues and the tissue-class inventory; scale bars, 200 µm. **B.** Class-balanced accuracy and macro-averaged precision, recall, and F1 across segmentation models. The highest mean score for each metric is shown in blue; other mean scores are orange, and gray markers indicate individual runs. **C.** Per-class precision (blue) and recall (orange) for TiSAM (UNI2-h) versus DeepLabV3+. Dashed diagonals indicate equality; axes begin at 30%. Labels identify the three largest gains (green) and deficits (red) for each metric. **D.** Per-class win counts and mean signed differences (TiSAM minus DeepLabV3+), with positive differences highlighted in green. All metrics are computed from class-balanced test confusion matrices containing 15,000 pixels per class.

Interestingly, SAM-Path and SAM2-Path showed a marked decrease in performance on MegaMouse relative to the publicly available datasets **(Fig. 2B-E**, **Fig. 3B)**. We hypothesize that this difference partly reflects the distinct annotation strategy used for MegaMouse. Unlike most public benchmarks, which provide exhaustively annotated ROIs, MegaMouse is made of smaller annotated regions that may include incomplete objects and are subsequently augmented and assembled into training tiles **(Fig. 3A, Fig. S7B)**. Such annotations may be challenging for methods trained with Dice-based objectives like SAM-Path and SAM2-Path, which explicitly optimize region-level overlap between predictions and ground-truth masks. In contrast, TiSAM uses median-frequency weighted cross-entropy, which provides a pixel-wise training objective and showed greater robustness across the heterogeneous annotation schemes evaluated here.

PathSegmentor showed particularly poor performance on MegaMouse, most notably in its out-of-the-box configuration. In contrast, fine-tuning substantially improved performance, bringing it closer to the levels observed on the public datasets **(Fig. 2B-E**, **Fig. 3B)**. This discrepancy may reflect the lack of animal histology data in its pre-training and development, which could limit its ability to generalize to the morphology and staining characteristics encountered in MegaMouse.

### TiSAM maintains robust performance in tasks with limited training data

To evaluate the performance of TiSAM in a multiclass segmentation task with limited training data, we developed a tissue segmentation training dataset consisting of diverse, high-quality, clinically derived human pancreas histology **(Fig. 4A).** The dataset comprised more than 12,000 expert-informed annotations across 50 WSIs from the Johns Hopkins Hospital (JHH) pathology archive, with annotations of major pancreatic structures: acini, islets of Langerhans, ducts, adipose tissue, collagen, smooth muscle, nerves, endothelium and immune aggregates, as well as the neoplastic precursor lesion pancreatic intraepithelial neoplasia (PanIN) **(Fig. 4A)**. The WSIs encompass variability in staining intensity and tissue preservation, including differences in staining protocols, fading, folding and tissue damage. To assess generalization beyond the training cohort, we established an independent test set of pancreatic histology from the JHH and public histology from GTEx^39^ and TCGA^40^.

**Figure 4:**
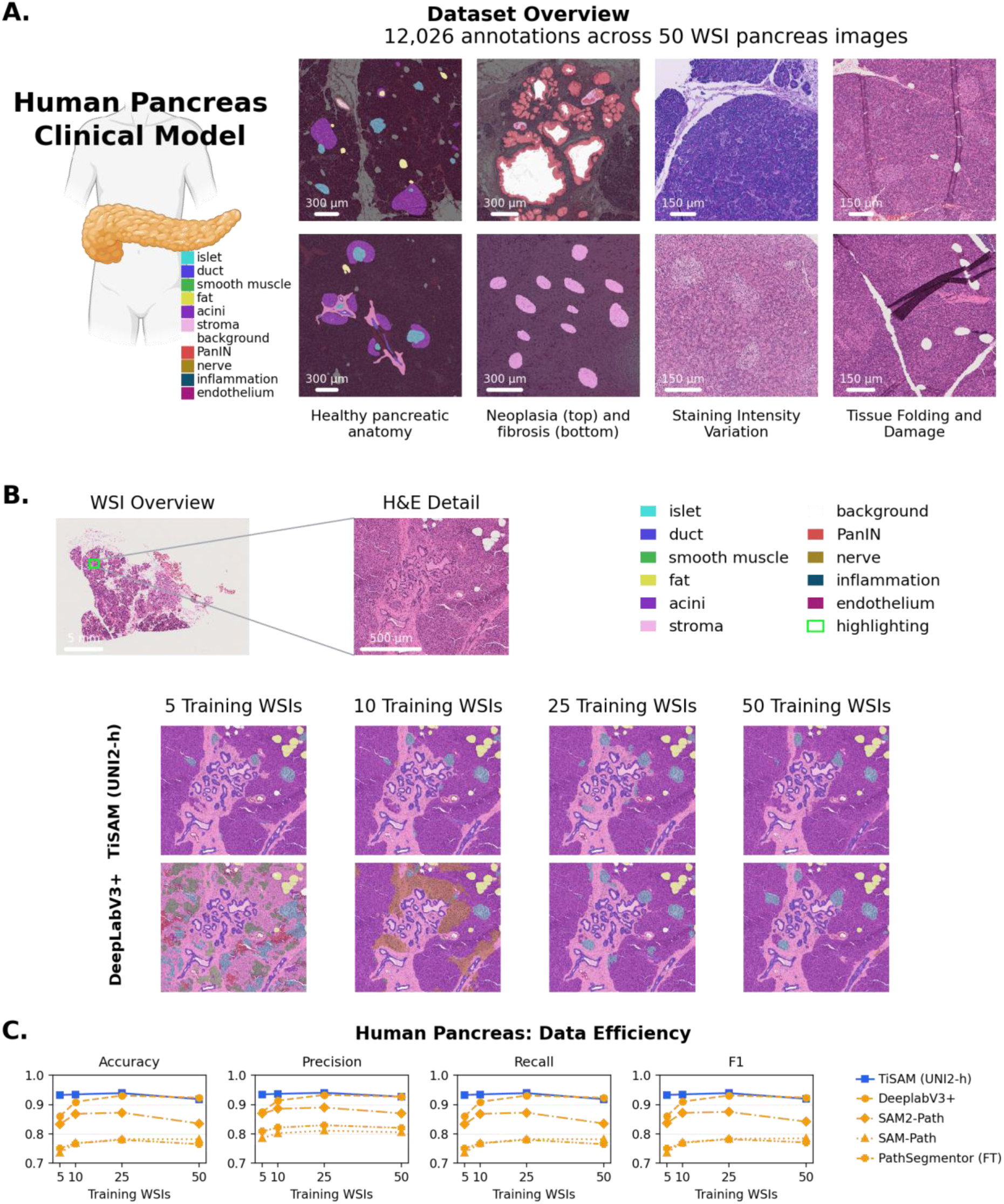
Human pancreas dataset overview and data efficiency. **A.** Representative annotated anatomy, including regions with neoplasia and fibrosis, staining variation, and tissue artifacts. **B.** Sample whole-slide image from the testing dataset, with TiSAM (UNI2-h) and DeepLabV3+ predictions for 5, 10, 25, and 50 training WSIs. **C.** Test accuracy and macro-averaged precision, recall, and F1 for the TiSAM (UNI2-h) and other models. All models were trained on the same pancreatic training subset at each training set size.

Clinical histology encompasses substantial variation in tissue preservation and cellular morphology, histological stain, and image quality. We hypothesized that the foundation-model representations incorporated into TiSAM would enable robust segmentation with fewer training annotations compared to existing architectures. To test this, we trained TiSAM and the previously mentioned model architectures using annotations from 5, 10, 25 and 50 WSIs

(corresponding to 2,218; 3,783; 8,458, and 12,026 annotations) from the human pancreas dataset and evaluated accuracy, precision, recall and F1 **(Fig. 4B, C)**. With only 5 or 10 training WSIs, TiSAM achieved macro-F1 scores of 93.3% and 93.5%, respectively, compared with 86.1% and 91.0% for DeepLabV3+, corresponding to advantages of 7.2 and 2.5 percentage points **(Fig. 4C)**. Consistent with the quantitative results, qualitative review showed that TiSAM correctly segmented complex regions even for limited training datasets, whereas the DeepLabV3+ model suffered from extensive misclassification when trained using 5 or 10 WSIs **(Fig. 4B)**. As the number of training WSIs increased from 5 to 25, DeepLabV3+ performance improved and approached that of TiSAM. Both models achieved their highest macro-F1 scores at 25 training WSIs, followed by lower scores at 50 WSIs, where DeepLabV3+ slightly exceeded TiSAM (92.4% vs. 92.0%) **(Fig. 4C)**.

DeepLabV3+ showed the greatest dependence on training set size, with substantial performance gains as the number of training WSIs increased from 5 to 25. The ViT-based architectures also benefited from increased training data, although none matched TiSAM performance across the evaluated training set sizes. PathSegmentor was the lowest performing architecture overall, including following fine-tuning on the pancreas training data **(Fig. 4C)**. Together, these results indicate that TiSAM can achieve strong segmentation performance with substantially fewer training annotations than conventional and alternative foundation-model-based architectures.

### TiSAM exhibits robust cross-scanner generalization

A major barrier to deploying AI models across institutions is performance degradation on out-of-distribution data^41,42^. One major contributor to this in the digital pathology space is variability in image color and texture introduced by cross-scanner variability. We hypothesized that the robust dual encoders in TiSAM would lead to improved cross-scanner performance compared to existing models such as DeepLabV3+ **(Fig. 5, Figs. S5 and S6)**.

**Figure 5:**
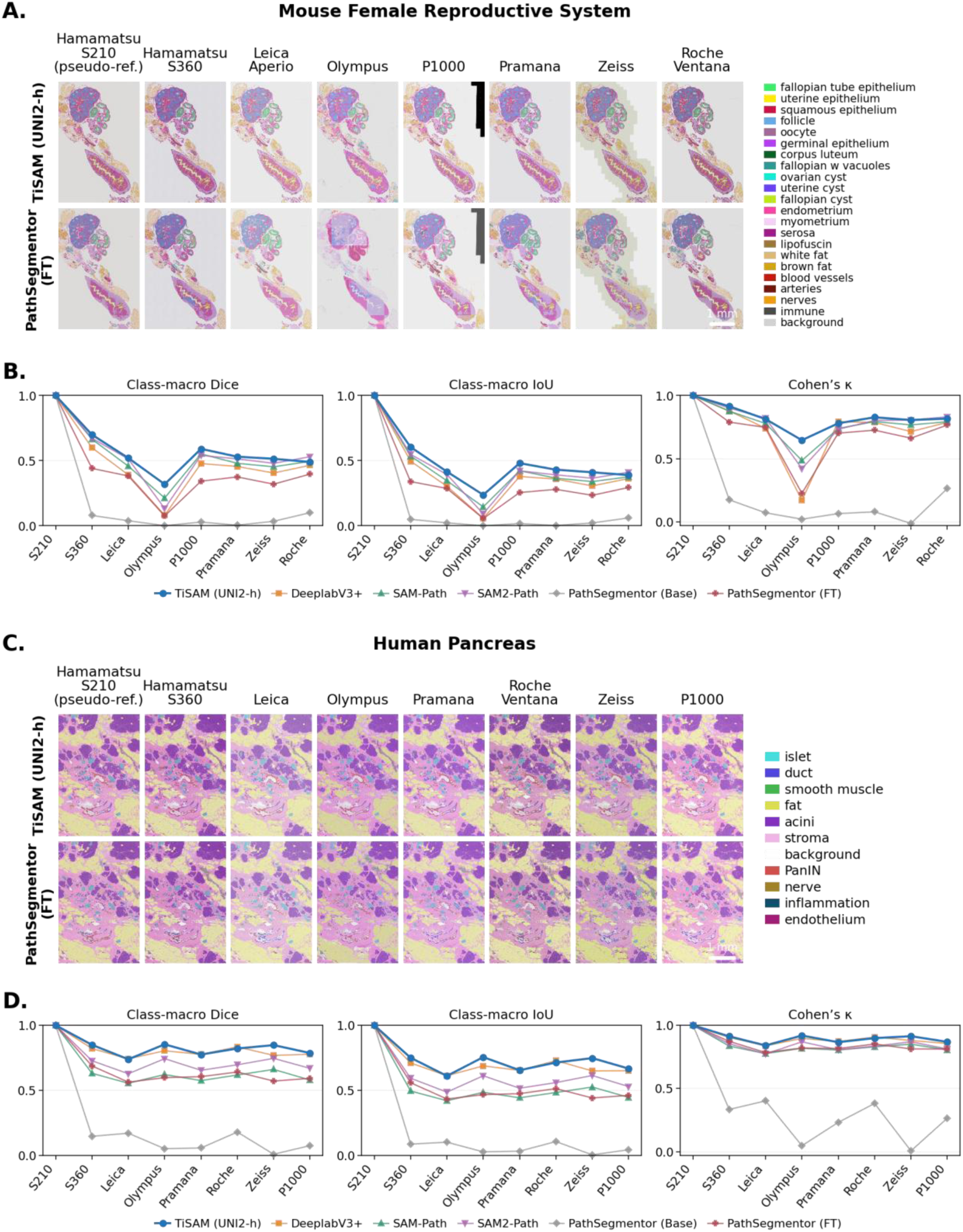
Cross-scanner prediction consistency. A, **C.** TiSAM (UNI2-h) and PathSegmentor (FT) predictions across eight scanners for matched (A) mouse female reproductive system and (C) human pancreas specimens. **B, D.** Macro Dice, Macro IoU, and Cohen’s kappa relative to each model’s Hamamatsu S210 prediction (fixed to 1). Each model is represented by a different color as shown in the legend. Metrics measure similarity to the segmentation of the Hamamatsu S210 scan, not manual annotation-based accuracy.

We first compared the cross-scanner generalization of TiSAM against the other models in the healthy female mouse reproductive system **(Fig. 5A, Fig. S6)**. The training dataset consisted of 47,738 annotations of 22 distinct tissue classes on a total of 156 WSIs digitized using a Hamamatsu NanoZoomer S210 scanner. We evaluated the performance of the models on a single tissue section serially scanned on eight scanners: Hamamatsu NanoZoomer S210, Hamamatsu NanoZoomer S360, Leica Aperio GT180, Olympus Slideview VS200, Pannoramic 1000 Digital Scanner, Pramana Spectral HT-2, ZEISS Axioscan 7, and Roche Ventana DP 600, using each model’s segmentation of the Hamamatsu S210 image as its pseudo-reference. TiSAM demonstrated superior overall cross-scanner consistency, achieving the highest macro-Dice on six of the seven non-reference scanners and outperforming competing models on most scanner-specific comparisons **(Fig. 5B)**. Most models showed their highest concordance on Hamamatsu S360 and their lowest on Olympus. TiSAM’s largest advantage over the nearest competitor was observed on Olympus, where it exceeded SAM-Path by 10.6 p.p. in macro-Dice and 16.1 p.p. in Cohen’s kappa. SAM2-Path was the closest comparator overall and led both metrics on Roche Ventana, where SAM-Path also slightly exceeded TiSAM in macro-Dice. DeepLabV3+ surpassed TiSAM in kappa on Pannoramic 1000. PathSeg (FT) substantially outperformed PathSeg (Base), but both remained below TiSAM across all scanners.

To confirm our observations, we repeated this experiment using the 50-WSI clinically derived human pancreatic tissue described above **(Fig. 5C, Fig. S5)**. TiSAM again showed higher overall concordance than PathSegmentor, although the differences were smaller than those observed in the mouse reproductive system **(Fig. 5D)**. Across scanners, TiSAM’s macro-Dice differences ranged from −1.3 to +7.9 p.p. relative to DeepLabV3+, +10.2 to +12.5 p.p. relative to SAM2-Path, +18.4 to +23.2 p.p. relative to SAM-Path, +16.2 to +27.6 p.p. relative to PathSeg (FT), and +56.8 to +83.8 p.p. relative to PathSeg (Base), making DeepLabV3+ the closest comparator.

All models showed substantially higher concordance on the human pancreas dataset than on the mouse reproductive system dataset, with mean macro-Dice increasing from 0.523 to 0.810 for TiSAM and from 0.410 to 0.789 for DeepLabV3+. TiSAM’s advantage over DeepLabV3+ narrowed, while its advantages over SAM2-Path, SAM-Path, and PathSeg (FT) widened. These differences may reflect the smaller pancreatic label set, with 11 tissue classes versus 22 in the mouse dataset^12^. The 50 clinically derived pancreatic WSIs may also provide greater variation in tissue presentation during training, helping models accommodate scanner-dependent differences. Conversely, although the mouse model was trained using a larger number of annotations, they were derived from high-quality research-grade histology from only 12 mice, providing less variation in image appearance compared to the pancreas dataset.

Together, these results indicate that TiSAM provides more consistent cross-scanner segmentation than existing pathology tissue segmentation models.

## Discussion

Semantic segmentation of histological images offers an opportunity to extract quantitative information from tissue at a scale that is difficult to achieve through manual annotation alone. Such approaches can support biological questions ranging from tissue organization during development to the characterization of pathological changes in a broad array of diseases^43–46^. Importantly, their utility extends beyond quantifying overall tissue composition: segmentation can also provide a means of selectively identifying and isolating specific tissue regions for downstream molecular characterization. Despite this potential, existing approaches often require extensive expert annotation and can lose performance when applied to tissue with different morphological complexity, image quality or acquisition conditions.

We developed TiSAM to address these limitations by combining complementary representations from general-purpose and histopathology foundation models through a multi-scale fusion architecture. Rather than simply concatenating SAM3 and pathology features, TiSAM projects pathology features into the 256-dimensional SAM3 feature space and selectively integrates the two representations across multiple resolutions. A pixel decoder then aggregates these features, while a query-based mask decoder combines semantic and spatial information to generate high-resolution tissue masks. This design allows TiSAM to leverage the semantic information learned by pathology foundation models while preserving the spatial and boundary information provided by SAM, likely contributing to its improved performance over existing approaches. Across datasets, TiSAM consistently matched or exceeded existing architectures.

A central advantage of TiSAM was its ability to generalize across the types of variation encountered in practical histopathology applications. TiSAM maintained stronger performance as the number and diversity of tissue classes increased, generalized more effectively to complex clinical pathology specimens and retained segmentation performance when substantially fewer manually annotated images were available for training. TiSAM also showed great robustness to color variation introduced by different scanners. To enable more rigorous assessment of these properties, we introduce two publicly available datasets, the human pancreas dataset and MegaMouse, which address challenges that are underrepresented in existing segmentation benchmarks, including complex clinical tissue, extensive tissue diversity, and cross-scanner variation.

Beyond its performance, the modular design of TiSAM provides a complementary alternative to the current trend towards zero-shot, universal segmentation models. These models have considerable potential to make histological analysis more accessible, but their training datasets are currently dominated by a limited range of human tissue types, restricting their ability to generalize to organs and tissue that are poorly represented or absent from their training data^15,18^. This limitation is particularly important for animal histology, which is largely absent from existing public segmentation datasets despite the commonality of animal experiments for studying development, disease, and treatment response. Building the large datasets required for truly universal models will therefore require substantial additional efforts in annotation and data curation. In the meantime, TiSAM offers a practical alternative by leveraging pretrained representations that can be efficiently adapted to specific biological or clinical applications with limited additional training. As larger and more diverse training resources become available, we envision lightweight task-adaptable frameworks such as TiSAM and universal segmentation models as complementary approaches for advancing scalable histological image analysis.

## Acknowledgements

The authors would like to thank Angelo DeMarzo, Tracy Jones, Kristen Lecksell, Alan Meeker, for the use of their brightfield slide scanners and to thank Michael Goggins for help with pathology image generation. The authors acknowledge the following sources of funding: Break Through Cancer; The Sol Goldman Pancreatic Cancer Research Center; P30CA006973.

## Disclosures

The authors declare no conflicts of interest.

## Data Availability

The training, validation, and test tiles for the MegaMouse and human pancreas datasets will be made available upon peer-reviewed publication.

## Code Availability

The code supporting this study are available in the following repository: https://github.com/Kiemen-Lab/tisam. An implementation of the model in a GUI is available in the following repository: https://github.com/Kiemen-Lab/CODAvision

## Materials and Methods

### TiSAM Architecture

TiSAM combines a frozen SAM3 image encoder with a frozen UNI2-h pathology encoder and a trainable semantic decoder **(Fig. 1A)**^17,21^. Given a 1024 × 1024-pixel RGB tile, the model predicts a semantic label at each pixel. Spatially gated fusion integrates the two encoders’ features, while a Mask2Former-style decoder converts the fused representations into dense segmentation outputs^24^.

The SAM3^17^ encoder produces a four-level feature pyramid with 256 channels per level. To incorporate domain-specific information, UNI2-h processes a 4 × 4 grid of image crops, each resized to 224 × 224 pixels. We retain its spatial patch tokens and arrange them according to their positions within the tile, producing a 1536-channel pathology feature map.

We fuse pathology features with the three coarser SAM3 pyramid levels. At each level, a separate 1 × 1 convolution projects the pathology features to 256 channels, followed by bilinear interpolation to match the spatial resolution. A sigmoid gate computed from the concatenated features assigns complementary weights to the SAM3 and pathology representations at each location. The fused representation is defined as follows:

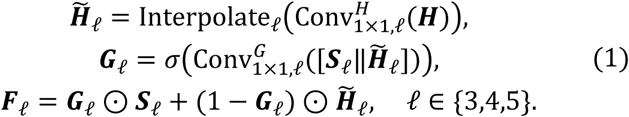

Here, ***S***_l_ and ***H*** denote the SAM3 and UNI2-h feature maps, respectively; *H̃*_l_ is the projected and resized pathology feature map. Superscripts *H* and *G* distinguish the projection and gate convolutions. The symbols ‖ and ⊙ denote channel-wise concatenation and elementwise multiplication. The gate ***G***_l_ is broadcast across channels. The batch dimension is omitted.

To preserve spatial detail in the highest-resolution SAM3 feature map, we retain the finest pyramid level without direct pathology fusion. A top-down pixel decoder combines the pyramid features and upsamples the resulting pixel embeddings to the input resolution before mask prediction.

A transformer decoder refines learnable queries through self-attention and cross-attention to the multi-scale image features. The human pancreas configuration uses 12 queries and six decoder layers, without additional SAM3 feature projection or image-side positional encoding. Pathology features are projected into the 256-dimensional SAM3 feature space for fusion, while SAM3 features are used without an additional projection. Each refined query produces class scores through a linear head and a mask embedding through a three-layer MLP. The query-specific mask logit at each pixel is the dot product between the mask embedding and the pixel embedding:

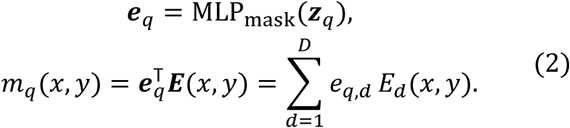

Here, ***z****_q_* is the final decoder representation of query *q*, ***e****_q_* ∈ ℝ*^D^* is its mask embedding, and ***E***(*x*, *y*) ∈ ℝ*^D^* is the pixel embedding, with *D* = 256.

We apply softmax to each query’s class scores and use these weights to sum the raw mask logits across queries, yielding the semantic logit for each class:

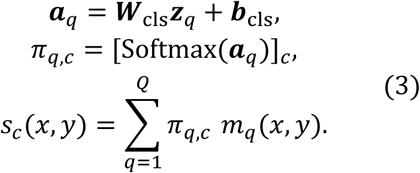

Here, ***W***_cls_ and ***b***_cls_ are the linear classification head parameters, *Q* is the number of queries, and *C* is the number of output classes, including synthetic void. The class probabilities *π_q_*_,*c*_ are normalized across classes for each query.

Queries therefore have no fixed class assignments, and training requires no Hungarian matching. Both encoders remain frozen while the fusion module, pixel decoder, query embeddings, transformer decoder, and prediction heads are trained.

### Datasets and Preprocessing

We evaluated TiSAM on three internal datasets and four public histopathology benchmarks. The human pancreas dataset contains 12,026 manual annotations on 50 WSIs across 11 classes: islet, duct, smooth muscle, fat, acini, stroma, background, PanIN, nerve, inflammation, and endothelium. The mouse female reproductive system dataset contains 47,738 manual annotations on 156 WSIs across 22 classes representing reproductive structures, supporting tissues, and background. MegaMouse combines a subset of annotations from the mouse female reproductive system dataset with samples from other mouse organ projects, comprising 36,833 manual annotations on 120 WSIs across 52 classes. These classes include shared tissue structures and organ-specific structures from 12 organs: lung, pancreas, skin, kidney, liver, bladder, heart, colon, vagina, uterus, fallopian tube, and ovary. Class counts include annotated background but exclude synthetic void.

The public benchmarks cover breast cancer, lung cancer, and melanoma. BCSS contains 151 annotated breast cancer images, divided into 85 training, 21 validation, and 45 test patients while preserving the official held-out test sites^26^. Following the original study, we grouped annotations into five classes: tumor, stroma, inflammatory infiltrates, necrosis, and other. Preparation yielded 1,541 training and 420 validation tiles, with test images retained at their original dimensions.

IGNITE H&E contains annotated non-small-cell lung cancer regions with 15 foreground classes and a background class. We used the official fold-0 split of 215 training and 54 validation regions, yielding 770 and 188 tiles, respectively, and evaluated on 139 publicly labeled test regions^27^. Training and validation were split at the region level and could share patients, whereas test patients were separate from both. Test images retained the supplied surrounding context, with unannotated pixels excluded from supervision and evaluation.

We used 300 annotated melanoma regions of interest (ROIs) from PUMA, excluding the 10 ROIs designated as the challenge’s preliminary test set. We divided the 206 training ROIs into 164 training and 42 validation ROIs and retained the 94 final test ROIs for evaluation, with equal representation of primary and metastatic melanoma in each split^28^. Each region represents a separate case and was supplied at 1,024 × 1,024 pixels, requiring no additional tiling or resizing during dataset preparation. Region identifiers served as case surrogates because patient and slide identifiers were unavailable. The labels comprise five tissue classes, stroma, blood vessel, tumor, epidermis, and necrosis, plus white background. Unlabeled pixels were ignored, and evaluation used the test partition.

BEETLE provides breast cancer annotations for four classes: non-invasive epithelium, invasive epithelium, necrosis, and other^29^. We used the official patient-level fold 0 for validation and the remaining four folds for training, yielding 34,648 packed training tiles and 10,324 source-aligned validation tiles. Test predictions for 170 images were submitted to the BEETLE challenge platform for scoring against hidden ground-truth labels.

Across datasets, training and validation inputs measured 1,024 × 1,024 pixels. Preparation included extracting image windows or assembling annotated regions into tiles, as required by each dataset. For assembled tiles, areas outside the extracted regions were filled with black pixels and assigned a synthetic void label, distinct from annotated background representing non-tissue whitespace **(Fig. S7)**. Models using SAM3 require interpolating input tiles to 1,008 × 1,008 pixels before feature extraction. Evaluation on images retained at their original dimensions used window extraction and prediction fusion to reconstruct masks within the original image bounds.

### Model Training

TiSAM and its variants, DeepLabV3+, HRNetV2-W18, SegFormer-B1, U-Net, and nnU-Net2D were trained using our PyTorch training pipeline for data loading, augmentation, optimization, and validation. These models used median-frequency weighted cross-entropy, with class weighting adopted from the CODA DeepLab implementation^9,47^. Comparisons of focal-plus-Dice loss and median-frequency weighted cross-entropy are provided for the mouse female reproductive system, MegaMouse, and human pancreas datasets in **Figs. S2-S4**, respectively. Standard TiSAM training kept the pretrained SAM3 and histopathology encoders frozen while optimizing the fusion and decoding components. Within this pipeline, DeepLabV3+ and U-Net used our code implementations. HRNetV2-W18 used a timm backbone with our segmentation head, SegFormer-B1 used Hugging Face Transformers, and nnU-Net2D used a fixed 2D PlainConvUNet from dynamic-network-architectures rather than the complete nnU-Net automated planning and training workflow.

SAM-Path and SAM2-Path were trained using the official implementations^22,23^. Their pretrained encoders remained frozen, while model-specific segmentation components were trained using the native objectives and optimization schedules. PathSegmentor (Base) used the authors’ released checkpoint without additional training^18^. PathSegmentor (FT) was fine-tuned from the released checkpoint using the official implementation and native training objectives. The image and language encoders remained frozen, while the pixel and transformer decoders were trained. Optimization settings, initialization, trainable components, training objectives, and checkpoint-selection rules are summarized in Tables S1 and S2.

### Evaluation

Segmentation performance was measured using accuracy and macro-averaged precision, recall, and F1. Internal datasets were evaluated using class-balanced confusion matrices, with each present annotated class sampled to the pixel count of the rarest class using a fixed seed. Synthetic void was excluded from macro-averaged metrics.

Public benchmarks were evaluated using full aggregate confusion matrices without class-balanced subsampling. Some tissue classes contained very few annotated pixels, so balancing all classes to the rarest class would discard most available pixels and make performance estimates sensitive to the sampled subset. We therefore retained all evaluated pixels and used macro-averaged precision, recall, and F1 to give each evaluated tissue class equal weight.

Where three runs were available, results were summarized as means and sample standard deviations across seeds 17, 42, and 137. Checkpoints in the controlled public benchmarks were selected using validation, without test-based checkpoint or seed selection. BEETLE test predictions were evaluated through the external platform using one validation-selected submission per model. We reported its official overall Dice and the unweighted mean of the four reported class Dice scores, including Other.

Cross-scanner consistency was assessed on two matched specimens, one human pancreas and one mouse female reproductive system specimen, each acquired with eight scanners and excluded from training. Each model’s Hamamatsu S210 prediction served as its pseudo-reference. Metrics were computed over the full available common support of aligned masks, independently of display crops. Pixel agreement and Cohen’s kappa included all pixels. Macro Dice and IoU included classes present in either mask, excluding synthetic void but retaining annotated background. These measurements quantify prediction consistency across scanner acquisitions rather than accuracy against manual annotations. For both the mouse female reproductive system and human pancreas cross-scanner experiments, all dataset-specific training and fine-tuning used images acquired exclusively with the Hamamatsu NanoZoomer S210 scanner. PathSegmentor (Base) used the released checkpoint without additional training.

## Supplementary Materials

**Figure S1:**
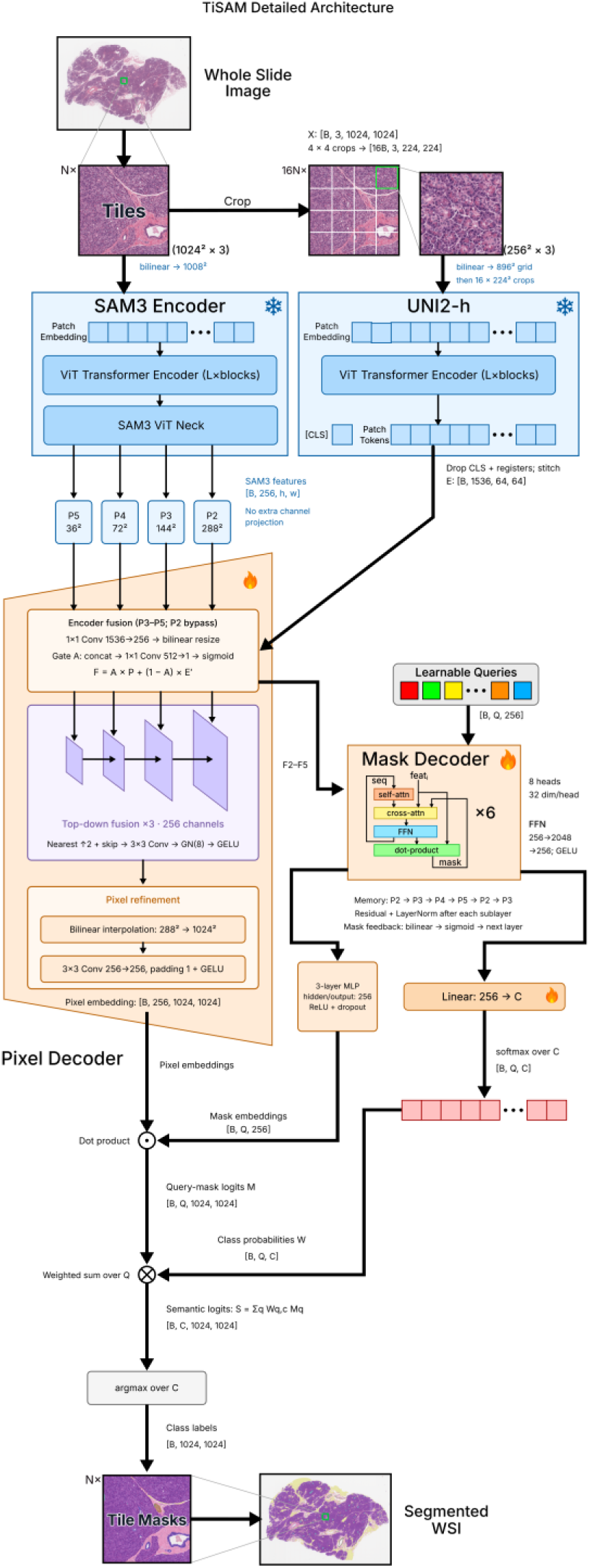
Detailed TiSAM architecture. A 1,024 × 1,024-pixel tissue tile is processed by frozen SAM3 and UNI2-h encoders. SAM3 receives a resized 1,008 × 1,008-pixel input, whereas UNI2-h encodes a grid of pathology crops. Gated fusion combines multiscale features, which are refined by the pixel decoder. Learned queries pass through the transformer decoder to produce class scores and mask embeddings. Their combination yields the semantic segmentation. Snowflakes mark frozen components.

**Figure S2:**
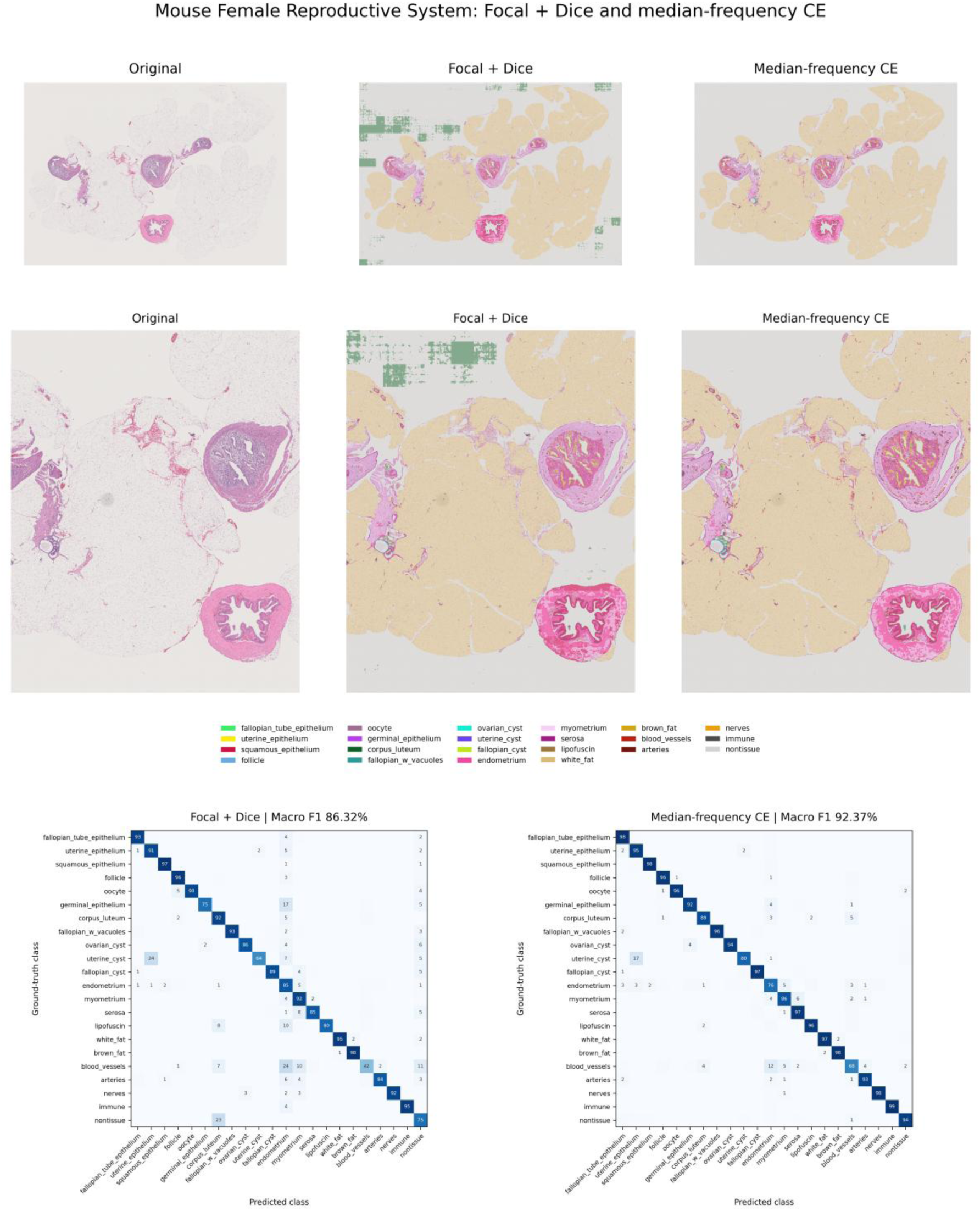
Mouse Female Reproductive System loss comparison. TiSAM (UNI2-h) predictions using focal-plus-Dice loss or median-frequency weighted cross-entropy. Rows show whole-slide overlays, a matched enlarged region of interest, and class-balanced test confusion matrices across 22 classes. Matrices are row-normalized, so diagonal entries indicate recall.

**Figure S3:**
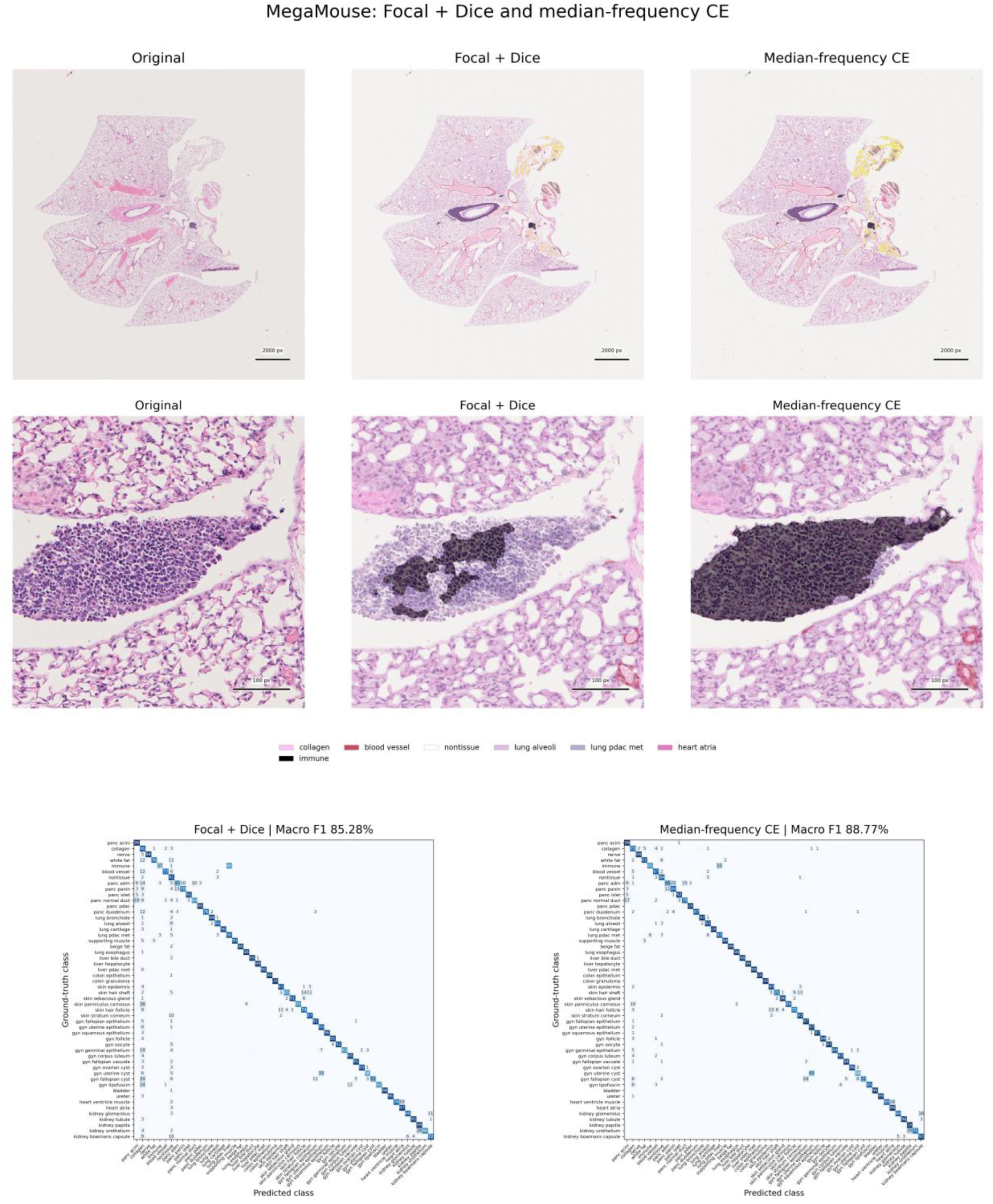
MegaMouse loss comparison. TiSAM (UNI2-h) predictions using focal-plus-Dice loss or median-frequency weighted cross-entropy. Rows show whole-slide overlays, a selected region of interest illustrating reduced nontissue overprediction, and class-balanced test confusion matrices across 52 classes. Matrices are row-normalized, so diagonal entries indicate recall.

**Figure S4:**
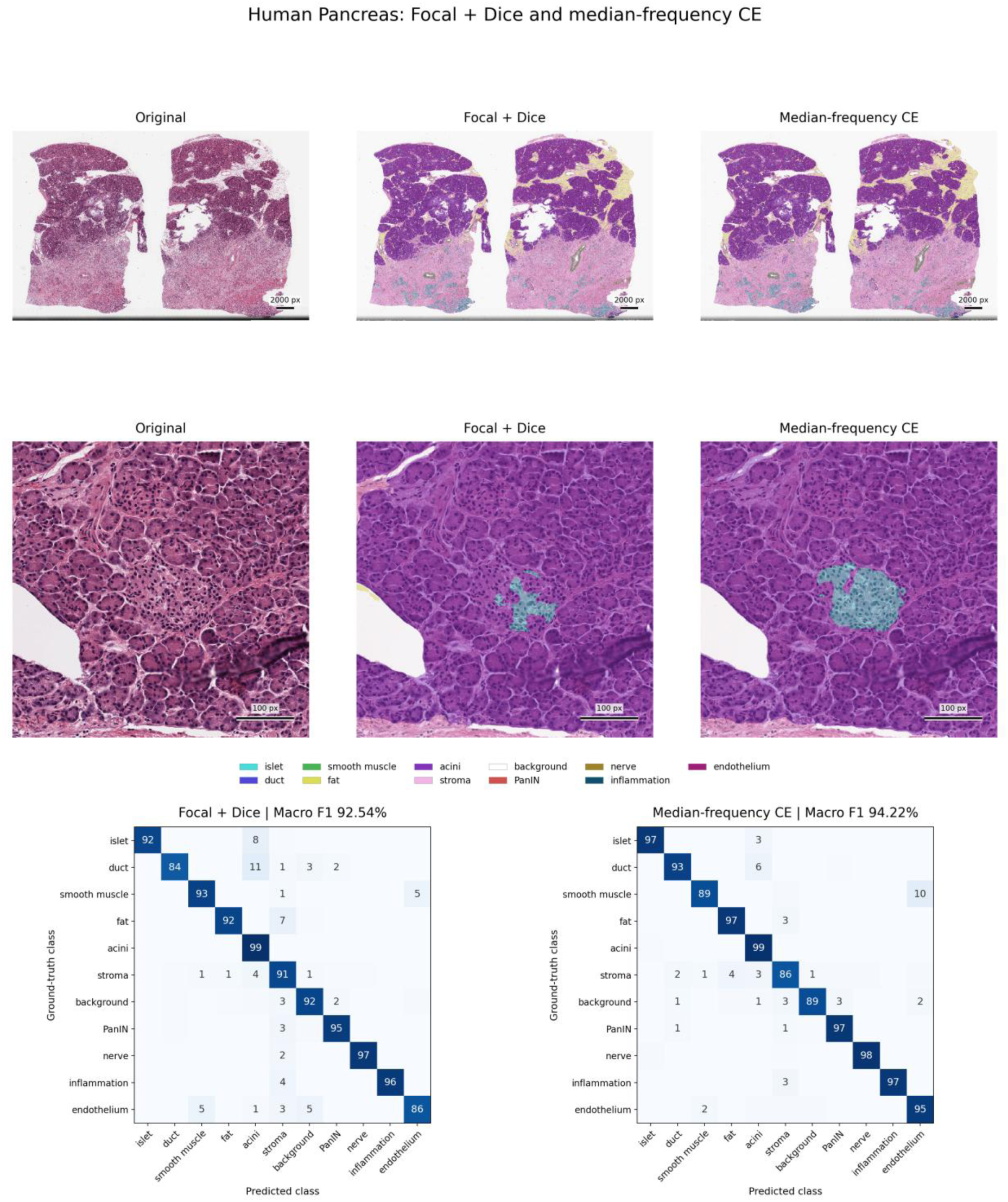
Human Pancreas loss comparison. TiSAM (UNI2-h) predictions using focal-plus-Dice loss or median-frequency weighted cross-entropy. Rows show whole-slide overlays, a selected region of interest illustrating improved islet delineation, and class-balanced test confusion matrices across 11 classes. Matrices are row-normalized, so diagonal entries indicate recall.

**Figure S5:**
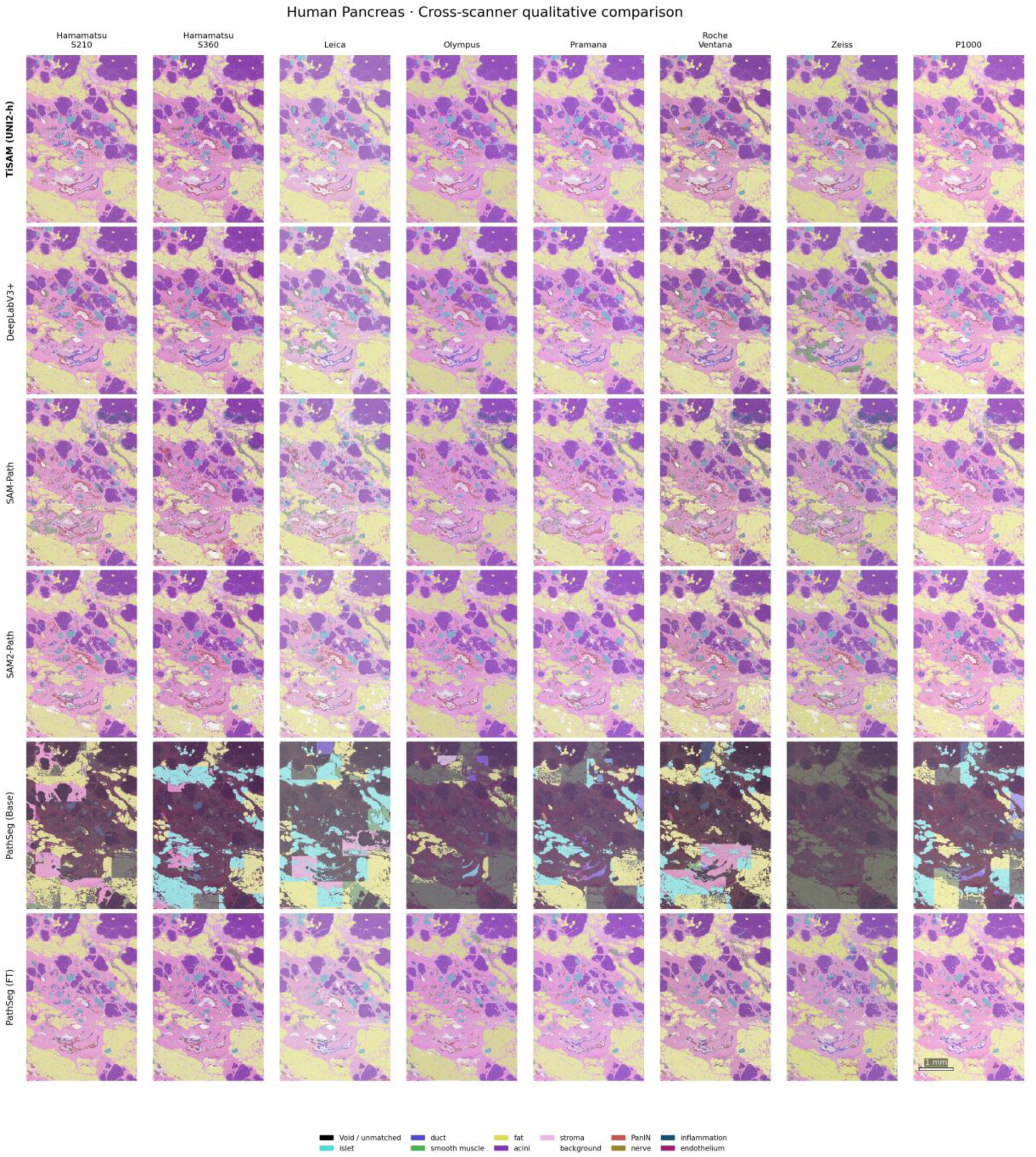
Human Pancreas cross-scanner comparison. Matched, registered regions of interest illustrate prediction consistency across eight scanners. Rows show TiSAM (UNI2-h), DeepLabV3+, SAM-Path, SAM2-Path, PathSegmentor (Base), and PathSegmentor (FT). Columns indicate scanners. PathSegmentor is abbreviated as PathSeg in the figure. Tissue colors are consistent across panels and are identified in the legend.

**Figure S6:**
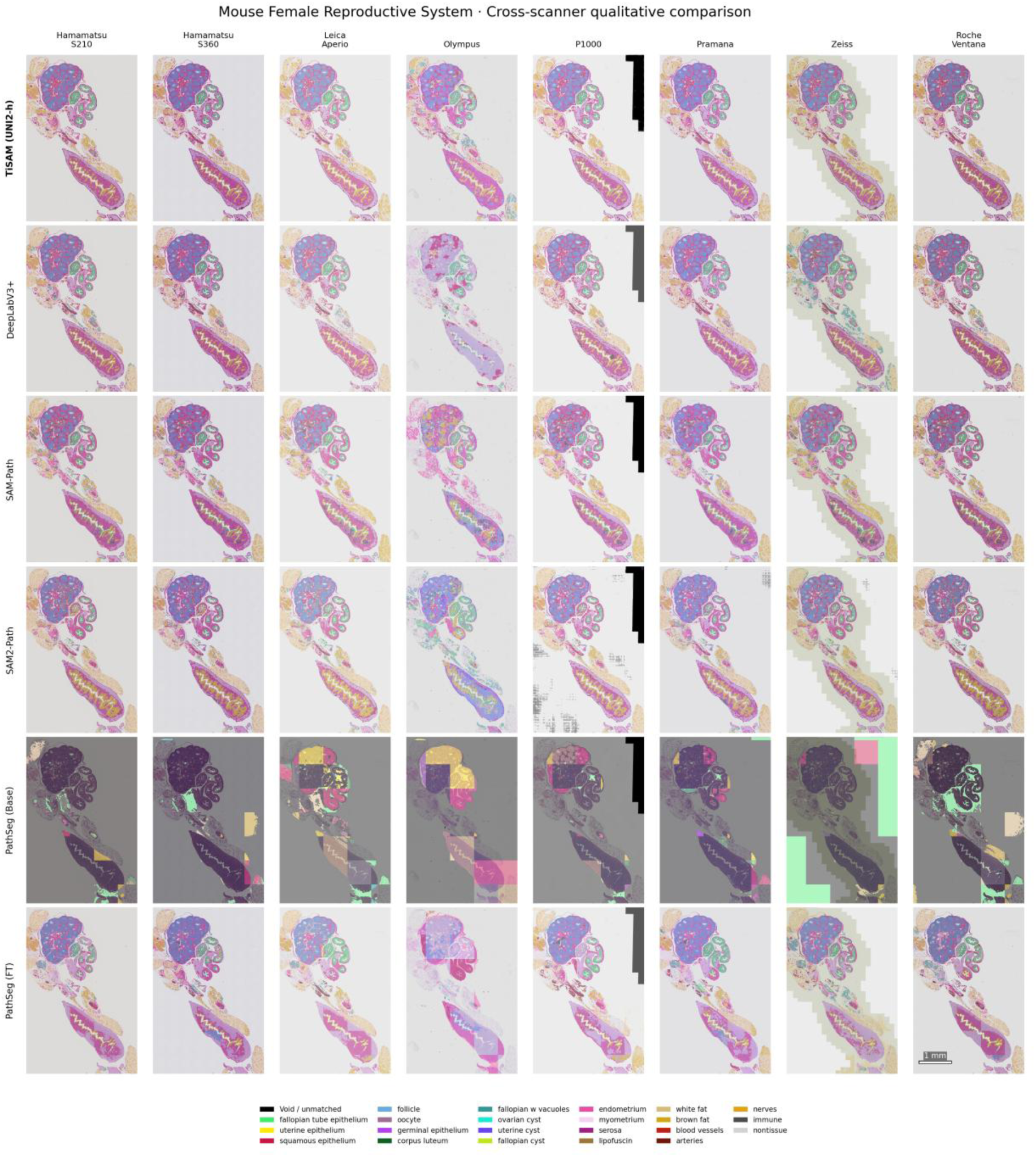
Mouse Female Reproductive System cross-scanner comparison. Matched, registered regions of interest illustrate prediction consistency across eight scanners. Rows show TiSAM (UNI2-h), DeepLabV3+, SAM-Path, SAM2-Path, PathSegmentor (Base), and PathSegmentor (FT). Columns indicate scanners. Dataset-specific trained models are shown alongside the released PathSegmentor (Base) reference. PathSegmentor is abbreviated as PathSeg in the figure. Tissue colors are identified in the legend.

**Figure S7:**
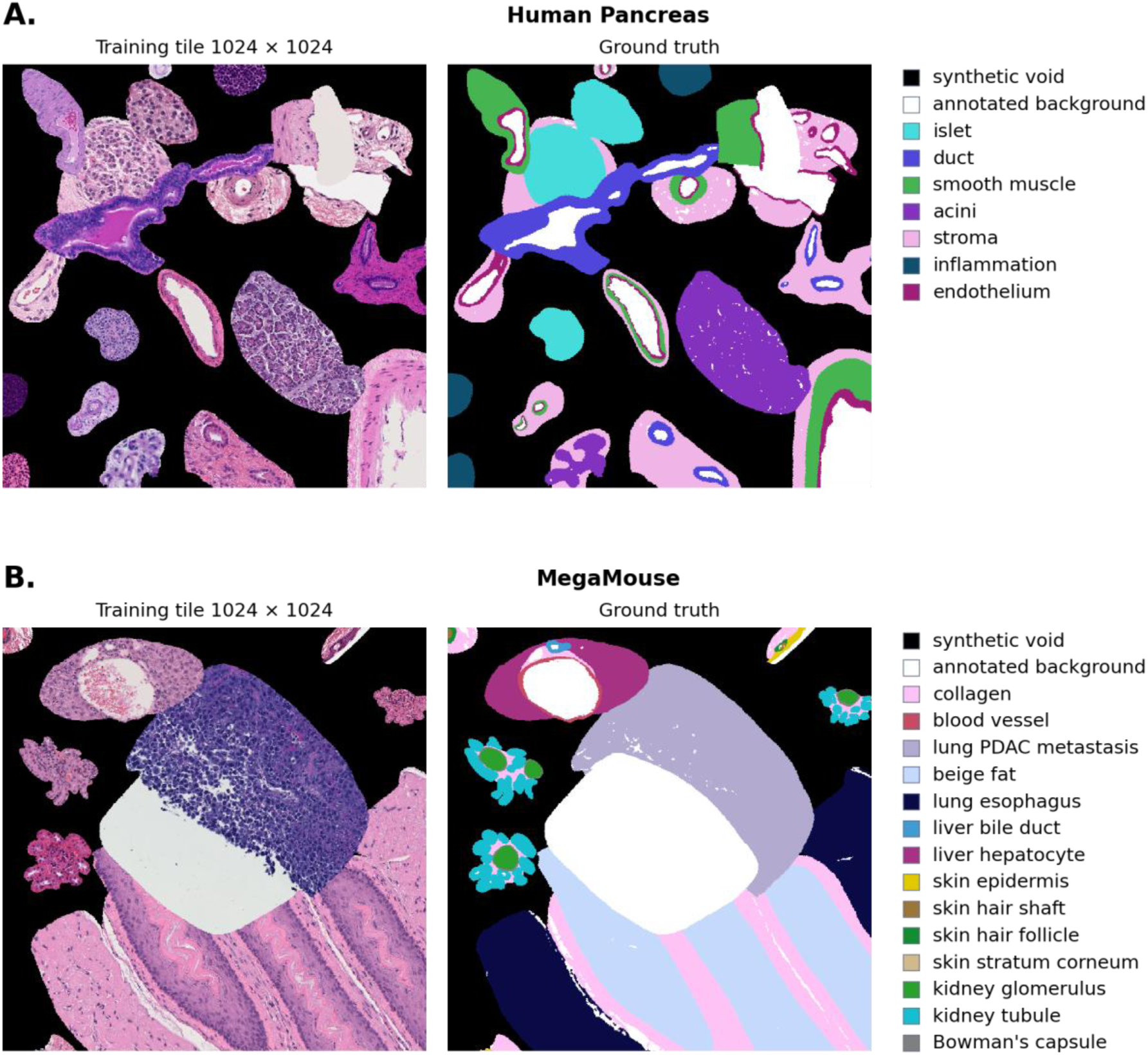
Examples of assembled training tiles and corresponding annotation masks. A, **B.** Examples from the human pancreas and MegaMouse datasets, respectively. Each row shows an H&E training tile (left) and its aligned semantic label mask (right). Annotated regions are assembled into a common tile, with spaces outside the extracted regions filled with black and assigned the synthetic void label. White regions in the masks represent annotated background, corresponding to non-tissue whitespace within the extracted regions, and are distinct from synthetic void. Other colors indicate tissue classes according to the accompanying legends.

**Figure S8:**
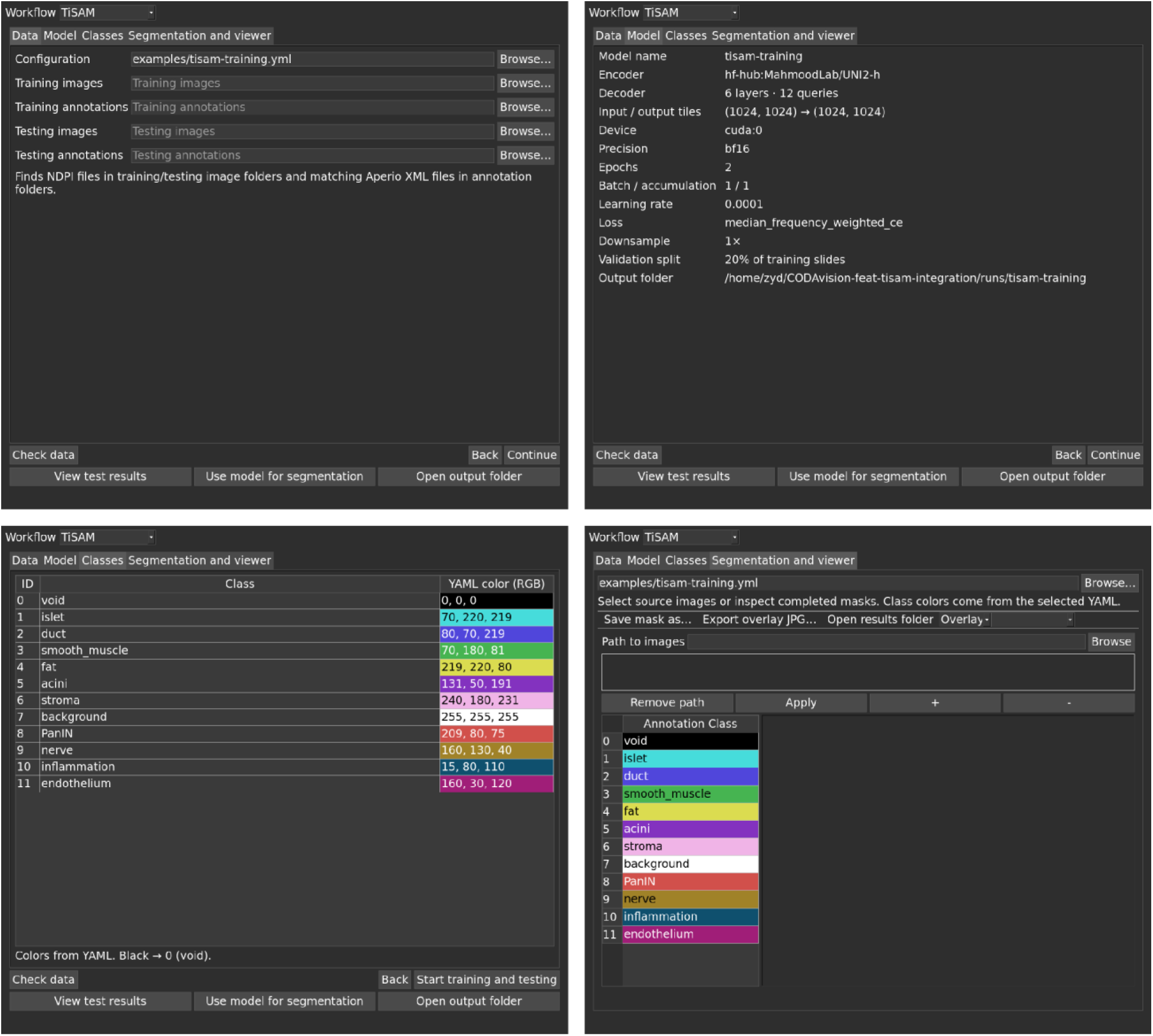
TiSAM integration in CODAvision. Screenshots of the CODAvision graphical user interface showing the integrated TiSAM workflow, including image and annotation selection, model and training settings, tissue class names and colors, and the segmentation viewer. The interface brings training, testing, and segmentation controls into a single application.

---

**Table S1:** Model training settings.

| Model and training context | Base LR, weight decay | Effective batch | Epoch limit | LR schedule | Early stopping and checkpoint selection |
| --- | --- | --- | --- | --- | --- |
| TiSAM (UNI2-h) | $5 \times 10^{-5}$, 0.1 | 16 | 200 | 5% linear warmup, linear decay | Loss ↓, patience 10, $\Delta = 0.001$ |
| TiSAM: final two UNI2-h blocks fine-tuned | $5 \times 10^{-5}$ decoder, $5 \times 10^{-6}$ encoder, 0.1 | 16 | 300 | 5% linear warmup, linear decay | Loss ↓, patience 10, $\Delta = 0.001$ |
| TiSAM (ResNet-50) | $5 \times 10^{-5}$, 0.1 | 16 | 200 | 5% linear warmup, linear decay | Loss ↓, patience 10, $\Delta = 0.001$ |
| TiSAM (UNI2-SEAL) | $5 \times 10^{-5}$, 0.1 | 16 | 200 | 5% linear warmup, linear decay | Loss ↓, patience 10, $\Delta = 0.001$ |
| DeepLabV3+ (ResNet-50) | $5 \times 10^{-4}$, 0.03 | 16 | 200 | 2% linear warmup, linear decay | mIoU ↑, patience 20, $\Delta = 0.0002$ |
| HRNetV2-W18 | $5 \times 10^{-5}$, 0.1 | 16 | 200 | 5% linear warmup, linear decay | mIoU ↑, patience 20, $\Delta = 0.0002$ |
| SegFormer-B1 | $5 \times 10^{-5}$, 0.1 | 16 | 200 | 5% linear warmup, linear decay | mIoU ↑, patience 20, $\Delta = 0.0002$ |
| U-Net | $5 \times 10^{-5}$, 0.1 | 16 | 200 | 5% linear warmup, linear decay | mIoU ↑, patience 20, $\Delta = 0.0002$ |
| nnU-Net2D | $5 \times 10^{-5}$, 0.1 | 16 | 200 | 5% linear warmup, linear decay | mIoU ↑, patience 20, $\Delta = 0.0002$ |
| SAM-Path | $10^{-4}$, 0.01 | 24 | 32 | 1-epoch warmup, $\times 0.1$ at epochs 25 and 29 | Fixed budget, best validation mIoU |
| SAM2-Path | $5 \times 10^{-4}$, 0.01 | 16 | 60 | 1-epoch warmup, $\times 0.1$ at epochs 25 and 29 | Fixed budget, best validation mIoU |
| PathSegmentor (FT) | $10^{-5}$, 0.05 | 1 | 20 | $\times 0.1$ at 88.889% and 96.296% of iterations | Fixed budget, best validation mIoU |

All trained models in this table use AdamW. Effective batch size includes gradient accumulation. LR denotes learning rate, $\Delta$ denotes the minimum improvement for early stopping, and mIoU denotes mean intersection over union across foreground classes. Loss and mIoU refer to validation measures. Arrows indicate the optimization direction. Patience is measured in epochs. Checkpoints follow the stated validation monitor. Epoch limits are maximum budgets, not necessarily completed epochs.
SAM-Path and SAM2-Path retain their official training recipes and fixed budgets. The earliest checkpoint is retained in a validation mIoU tie. The PathSegmentor (FT) settings shown follow the official 20-epoch training recipe. These settings apply to all three seeds across all datasets. The nominal ten warmup iterations use a factor of 1.0 and do not ramp the learning rate.
Benchmark comparisons use seeds 17, 42, and 137. Checkpoint selection uses validation data.

**Table S2:** Model implementation, initialization, and training objective.

| Model | Implementation / training pipeline | Initialization / trainable components | Objective |
| --- | --- | --- | --- |
| TiSAM (UNI2-h) | Ours | Pretrained SAM3 and UNI2-h frozen. Fusion and decoder trained. | Weighted CE |
| TiSAM: final two UNI2-h blocks fine-tuned | Ours | SAM3 and remaining UNI2-h blocks frozen. Fusion, decoder, and final two UNI2-h blocks trained. Encoder LR = $0.1 \times$ decoder LR. | Weighted CE |
| TiSAM (ResNet-50) | Ours, torchvision backbone | ImageNet ResNet-50. Full backbone and TiSAM decoder trained. | Weighted CE |
| TiSAM (UNI2-SEAL) | Ours | SAM3 and released UNI2-SEAL frozen. Fusion and decoder initialized from TiSAM (UNI2-h), training state reset. | Weighted CE |
| DeepLabV3+ (ResNet-50) | Ours, torchvision backbone | ImageNet ResNet-50. Full backbone and head trained. | Weighted CE |
| HRNetV2-W18 | timm backbone, our head / ours | ImageNet initialization. Full backbone and head trained. | Weighted CE |
| SegFormer-B1 | Hugging Face Transformers / ours | Random initialization. Full model trained. | Weighted CE |
| U-Net | Ours | Kaiming initialization. Full model trained. | Weighted CE |
| nnU-Net2D | Upstream PlainConvUNet / ours | Kaiming initialization. Full model trained without deep supervision. | Weighted CE |
| SAM-Path | Official implementation | Pretrained SAM ViT-B and HIPT encoders frozen. Dimensionality reduction module, class prompts, and mask decoder trained. | 0.25 focal + 0.75 Dice + 0.0625 IoU regression |
| SAM2-Path | Official implementation | Pretrained SAM2 Hiera-B+ and UNI encoders frozen. Fusion module, class prompts, and SAM-based mask decoder trained. | 0.25 focal + 0.75 Dice + 0.0625 IoU regression |
| PathSegmentor (FT) | Official implementation | Released checkpoint. Image and language encoders frozen. Pixel and transformer decoders trained. | Native classification, mask BCE and Dice, grounding, and spatial losses |

Ours denotes our implementation and training pipeline unless another implementation source is specified. TiSAM fusion and decoder include the fusion module, pixel decoder, query embeddings, transformer decoder, and prediction heads. TiSAM input tiles are $1,024 \times 1,024$ pixels, resized to $1,008 \times 1,008$ for SAM3. Weighted CE denotes median-frequency weighted cross-entropy, with weighting adopted from CODA DeepLab and class weights estimated from the training partition. Architecture libraries are dynamic-network-architectures 0.4.4 (nnU-Net2D), Transformers 5.9.0 (SegFormer-B1), and timm 1.0.24 (HRNetV2-W18). nnU-Net2D uses only the 2D PlainConvUNet architecture, without nnU-Net planning, preprocessing, or training.
SAM-Path and SAM2-Path retain their native objectives and budgets, with dataset-specific classes and ignored-label handling. PathSegmentor retains its official training entry point, optimizer, scheduler, loss weights, auxiliary supervision, and FP16 automatic mixed precision. Fine-tuning uses dataset-specific class names, 20 epochs, and gradient clipping at 5.0. Mask matching and loss reductions exclude unannotated void points. BCE denotes binary cross-entropy.

## Notes

### Competing Interest Statement

The authors have declared no competing interest.

